# T2R14 mediated antimicrobial responses through interactions with CFTR

**DOI:** 10.1101/2024.04.25.591203

**Authors:** Tejas Gupte, Nisha Singh, Vikram Bhatia, Kavisha Arora, Shayan Amiri, Paul Fernhyhough, Anjaparavanda P Naren, Shyamala Dakshinamurti, Prashen Chelikani

**Author notes:** **Corresponding author:** P. Chelikani, D319, Manitoba Chemosensory Biology Research Group and Department of Oral Biology, Dr. Gerald Niznick College of Dentistry, Rady Faculty of Health Sciences, University of Manitoba, 780 Bannatyne Avenue, Winnipeg, MB R3E 0W4, Canada.

## Abstract

Bitter taste receptors (T2Rs), are a subset of G protein-coupled receptors (GPCRs) that play a key role in responding to microbial presence at epithelial surfaces. In epithelia, the activities of ion channels and transporters, and of T2Rs, mutually affect each other. The normal function of one such anion channel, cystic fibrosis transmembrane conductance regulator (CFTR), is essential for the maintenance of healthy epithelia, not just in the respiratory but in the digestive and reproductive system as well. Based on evidence that T2R14 activity is affected upon mutations in *CFTR*, we explored the possibility that T2R14 and CFTR directly interact in cell membranes. The biophysical interaction between these proteins was mapped to specific regions of the CFTR, and was dependent on agonist stimulation of T2R14. Further, T2R14 was found to couple to Gαq, in addition to the canonical Gαi, in response to bacterial and fungal quorum sensing molecules. Whether the interaction with CFTR affects T2R14 driven responses to microbial signals is under investigation.

## INTRODUCTION

Epithelial surfaces form a physical barrier that prevents microbial access to host tissues. Microbes and microbial signals are detected by pattern recognition receptors on epithelial cells, which then coordinate innate immune responses. The innate immune response of epithelial cells to a bacterial challenge is illustrated by Toll-like receptors (TLRs) that recognize various pathogen associated molecular patterns (PAMPs) and respond by secreting cytokine (chemokines, interferons) and/or antimicrobial peptides (mucins and defensins) [1]. While TLRs are cell surface receptors, NOD-like receptors (NLRs) function as intracellular pattern recognition receptors (PRRs) [2]. Recently it has been recognized that a separate group of receptors may function in parallel to supplement the innate immune response [2]. Taste 2 receptors (T2Rs) are a collection of low affinity G protein- coupled receptors (GPCRs) that were originally discovered to detect bitter tastants upon exposure to these compounds in the taste buds [3, 4]. Subsequent studies show that T2Rs exhibit extra-oral expression in various tissues, and that they detect and respond to other non-gustatory signals as well [5, 6].

Bitter agonist-mediated T2R stimulation in human smooth muscle cells leads to bronchodilation [7], and vasoconstriction in the pulmonary circuit [8]. A specific bitter taste receptor, T2R38, was identified as the receptor for the quorum sensing molecule (QSM) – *N*-(3-oxododecanoyl)-L-homoserine lactone (AHL-C12)- from Gram negative bacteria such as *Pseudomonas aeruginosa*, in the human upper respiratory epithelium [9]. Acyl homoserine lactones (AHLs), of which AHL-C12 is member, are commonly occurring QSMs. AHL-stimulation of T2R38 led to NO secretion, an increase in ciliary frequency and enhanced mucociliary clearance. Subsequently, T2R4, T2R14, and T2R16 were identified for their ability to recognize bacterial quinolones [10]. The binding site for various AHLs and quinolones were dissected for T2R1, T2R4, T2R14 and T2R20 [11]. T2R14 also detects QSMs from Gram positive bacteria, including Competence Stimulating Peptides (CSP1) from *Streptococcus mutans* [12]. Recognition of various bacterial-derived signals by T2Rs elicits innate immune responses in terms of NO production and secretion, and secretion of human beta-defensin and interleukins [13]. Thus, bitter taste receptors may be considered a redundant PRR component that contribute to antimicrobial responses and pathogen clearance, particularly from the respiratory tract.

Signaling pathways downstream of agonist-stimulated T2Rs have been examined, and found to involve gustducin [3, 4] and Gαi2 [14, 15]. In parallel, agonist-stimulation of T2Rs precipitates changes in Ca^2+^ through Gβγ-mediated PLCβ [16], and Gαi activation reduces cAMP accumulation by inhibiting adenylate cyclase [17, 18]. Structural investigations suggest T2R46 precoupling to gustducin [19] as well as basal activity through T2R14-Gαi1 and T2R14-gustducin [20]. On the other hand, a recent study implicates Gαq in Gβγ-mediated activation of PLCβ [21]. These disparate observations about the G proteins and secondary messengers operating downstream of T2Rs suggest that signaling downstream of T2Rs would benefit from a detailed characterization.

Cystic fibrosis (CF) is a well-characterized genetic disorder caused by mutations in the *CFTR* gene which encodes the Cystic fibrosis transmembrane conductance regulator (CFTR) protein [22, 23]. CFTR is an anion channel that transports chloride [24, 25] and bicarbonate ions [26], and is gated by ATP hydrolysis [27], phosphorylation [28] and Calcium/Calmodulin binding [29]. People with CF (pwCF) present with various symptoms including production of abnormal mucus, impaired ciliary movement and compromised mucociliary clearance [30]. Together, these defects increase susceptibility to infection by opportunistic pathogens including the Gram-negative bacterium *P. aeruginosa,* and fungi like *Aspergillus fumigatus* and *Candida spp* [31, 32].

Recently, it has been reported that the loss of CFTR function results in diminished T2R-mediated innate immune responses [33]. However, the mechanism by which CFTR function affects T2R-mediated cellular responses has not been investigated. There is additional evidence attesting to bidirectional regulation of activities between ion channels, including CFTR, at one end and T2Rs at the other. Specifically, the bitter compound denatonium inhibits epithelial sodium channel ENaC [34], and an inhibitor of CFTR reduces the magnitude of response to denatonium. Bitter tasting compounds stimulate chloride secretion from rat ileum [35] and regulate potassium currents [36]. These observations led us to hypothesize that CFTR could interact directly with specific T2Rs, and that such a protein-protein interaction (PPI) could modulate signaling downstream of T2Rs. We have previously determined that T2R14 plays a significant role in eliciting responses to pathogens of significance in pwCF. Also, T2R14 is significantly overexpressed in various extra-oral tissues, including bronchial epithelial cells [20, 37]. Therefore, we specifically queried if CFTR interacts directly with T2R14 and how such an interaction could influence downstream signaling. Here we test the role of CFTR – T2R14 interaction on cellular response to bacterial (AHL-C12) and fungal (farnesol, tyrosol) QSMs.

Experiments were performed employing biophysical and pharmacological approaches, protein engineering tools and signal transduction assays. The results indicate that CFTR and T2R14 interact, however, the interactions are detected only in response to specific T2R14 agonists. These interactions can be attributed to either the N-terminus lasso motif or the NBD2 and C-terminus domain of CFTR. All the ligands tested T2R14 coupling to Gαq, in addition to the recognized Gαi cognate G protein. Efforts are ongoing to understand the ligand bias by different microbial QSMs on signaling through these pathways, and the effect of CFTR interaction on this signaling network.

## MATERIALS AND METHODS

### Compounds, stock solutions and buffers

Diphenhydramine (DPH), farnesol, tyrosol, *N*-(3-oxododecanoyl)-L-homoserine lactone (AHL-C12) and 6-methoxyflavanone were purchased from Sigma. Stock solutions of these compounds (DPH, 500 mM, farnesol 45 mM, tyrosol 145 mM, AHL-C12 166 mM, 6-methoxyflavanone 100 mM) were prepared in DMSO and stored at −20°C as single use aliquots. Compounds other than DPH were diluted from DMSO into ethanol, and finally into buffer A (below) for application to cells. The aqueous solutions were vigorously vortexed before use. Inorganic salts and buffers were purchased from Sigma. Buffer A containing 140 mM NaCl, 5 mM KCl, 2 mM CaCl_2_, 1 mM MgCl_2_, 10 mM HEPES (pH 7.4) mimicking extracellular ionic composition was prepared and filtered (0.45 µ syringe filter) immediately before use.

### Cell culture consumables and reagents

Cell culture media, supplements, dissociation reagents (Trypsin, Accutase) and phosphate buffered saline (PBS) were purchased from Invitrogen (Carlsbad, CA, USA) and Cedarlane, Canada. Monoclonal anti-HA (mouse IgG1, Invitrogen 26183) and goat anti-mouse Alexa488 (highly cross adsorbed, Invitrogen A32723) were purchased and stored in single use aliquots. Plasmid midiprep kit was purchased from Qiagen. NanoBRET NanoGlo detection system (Promega, N1662) and NanoGlo Live cell assay system (Promega, N2012) were used for NanoBRET and split Nanoluciferase complementation Systematic Protein Affinity Strength Modulation (SPASM) assays respectively. 96 well, white, solid, flat bottom polystyrene assay plates were purchased from Corning (Costar 3922) or Becton Dickinson (3296). For immunofluorescence, black plates with transparent bottom (Costar 3603) were used.

### Plasmids and molecular biology

EYFP-CFTR has been reported previously [38]. All other constructs were generated in a pcDNA3.1-Hygro(+) backbone. Cloning was performed by Genscript as per complete insert sequences provided, and plasmids were confirmed by sequencing before use. Common connecting sequences include the ER/K 10 nm linker used in the SPASM sensors derived from the myosin VI [39], and the ‘block’ domain that introduces rigidity derived from TNF receptor associated factor 2 (TRAF2) [40]. T2R14 was encoded with C-terminus Nanoluciferase for BRET. Control constructs for BRET include the pcDNA3.1-EYFP, pcDNA3.1-Nanoluciferase, pcDNA 3.1 EYFP-Nanoluciferase that encodes EYFP – GSS linker – Nanoluciferase and serves as the positive control for BRET validation, and EYFP – block – Nanolcuiferase which is the negative control for BRET [40]. For the SPASM sensors, T2R14 was tagged with 33 amino acids from the N-terminus of Rhodopsin to improve cell surface expression [41], and Hemagglutinin tag at the N-terminus for immunofluorescence detection. At the C-terminus, the SPASM module encoding the LgBiT of Nanoluciferase, 10 nm ER/K linker and SmBiT of Nanoluciferase were added. Finally, at the C-terminus of the construct, individual cytoplasmic domains of CFTR were added as follows N-terminus (1-80), NBD1+R-domain (351-850), R domain (595-860) and NBD2/C-terminus (1151-1480). For NanoBRET, T2R14 was tagged at the C-terminus with either Nanoluciferase or HaloTag. Each G-protein (Gαi2, Gαq or Gustducin) was developed in 4 versions – N-terminus Nanoluciferase, N-terminus HaloTag, C-terminus Nanoluciferase, C-terminus HaloTag. Plasmids were transformed into *E.coli*, using either DH5α or JM109 (*recA^-^, endA^-^,* purchased from Addgene) specifically to stabilize the SPASM sensor plasmids. The transformed bacterial clones were amplified in LB broth with Ampicillin (100 µg/mL) and plasmid DNA purified by standard midiprep procedure using Qiagen MidiPrep kit. DNA concentration and purity was estimated using a NanoDrop.

### Cell culture and Transfections

HEK293T cells (ATCC CRL-3216) were cultured in DMEM/Ham’s F-12 medium supplemented with 2.4 g/L sodium bicarbonate, 10% heat inactivated fetal bovine serum, 100 I.U /mL penicillin, 100 µg/mL streptomycin, on sterile cell culture treated plasticware, in 5% CO_2_ and appropriate humidity at 37°C. Cells were passaged upon reaching 90% confluence. For transfections, 0.3 x 10^6^ cells/mL were plated in 6-well (2 mL/well) / 12-well (1 mL/well) plates that had been coated *a priori* with poly-L-lysine (1 mg/mL), washed and dried while maintaining sterility. Following overnight adherence and growth, cells were transfected with desired plasmids using Lipofectamine 2000, and OptiMEM reduced serum medium, as per manufacturers’ instructions. Final concentrations were 1µg DNA and 2 µL Liopfectamine 2000 per mL of culture medium. For BRET, T2R14-Nanoluciferase and EYFP-CFTR were transfected at 500 ng each for the single concentration measurement. For the testing BRET at different acceptor:donor ratios, T2R14-Nanoluciferase was held constant at 250 ng, and EYFP-CFTR plasmid was varied. Donor alone control transfections were set-up simultaneously with the same amount of T2R14-Nanoluciferase DNA. Total DNA used in transfections was kept constant by addition of pcDNA3.1 empty vector. SPASM sensors had differing expression upon transfection. Sensor expression was monitored by immunofluorescence against the HA tag to determine cell surface expression, and DNA amounts optimized to get equivalent sensor expression. For assays with SPASM-sensor expressing cells the following DNA amounts were used, N-terminus 0.5 µg, NBD1+R 2 µg, R-domain 1.5 µg, NBD2/C-terminus 1 µg. Total DNA was adjusted to 2 µg as required by adding pcDNA3.1 empty vector. For NanoBRET assays, cells were transfected with 200 ng T2R14-Nanoluciferase combined with 1.8 µg of the complementary Gα-HaloTag. All transfections were allowed to proceed to 30 hours. Media was changed to maintain cell health at ∼24 hr after transfection.

At 30 hr after transfection, cells were washed with 1x PBS, detached into a single cell solution using Accutase which was quenched by adding the culture medium. Cell density was counted in a BioRad TC20 automated cell counter, and viable cell density was adjusted to 2.2 x 10^6^ cells/mL in OptiMEM reduced serum medium (no serum added). 90 µL of the cell suspension was pipetted into each well in a 96-well, poly-L-lysine coated plate to achieve 20,000 cells/well. Plates were centrifuged gently at 100x*g* for 30 seconds, and incubated at 37°C for 2 hr before the assay. For immunofluorescence assays, 60,000 cells were plated per well in a black, transparent plate. For NanoBRET assay, after adjusting cell density, the solution is split in half, with one half receiving Halo618 for HaloTag (acceptor) labeling and the other receiving DMSO (unlabelled acceptor is a donor alone control), as per manufacturer’s instructions.

### BRET (Bioluminescence Resonance Energy Transfer) and NanoBRET

Cells expressing T2R14-Nanoluciferase with or without EYFP-CFTR were stimulated with DPH (final concentration 2 mM, resuspended in buffer A) using 10 µL of 20 mM solution, or treated with buffer A containing 0.4% DMSO (to control for DMSO in the DPH stock solution). The plate was centrifuged at 100x*g* for 30 seconds, cells were incubated for 5 minutes, with orbital shaking in a SpectraMax iD5 plate reader equilibrated to 37°C. Subsequently 25 µL NanoBRET NanoGlo Substrate (diluted 1:1000 in buffer A) was added to each well and the plate was centrifuged as before, and returned to the plate reader. Following 30 seconds shaking, the luminescence signal and the resonance (EYFP) signal was measured at 447 nm and 535 nm respectively using appropriate bandpass filters and 500 ms integration time. BRET ratio and Net BRET were calculated as below

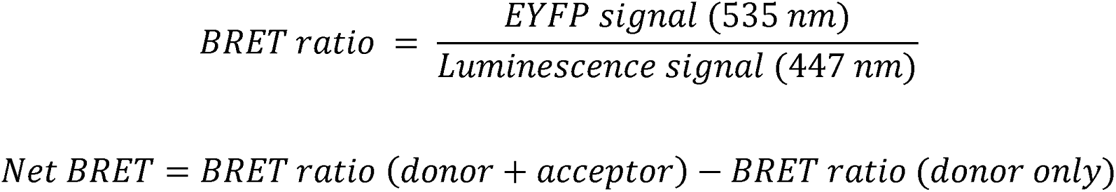

Compounds for NanoBRET were applied at the following final concentrations- DPH 2 mM, AHL-C12 200 µM, Farnesol 200 µM and Tyrosol 200 µM. AHL-C12 was diluted from DMSO stock into buffer A. AHL-C12, Farnesol and Tyrosol were diluted from DMSO stock into ethanol, and then into buffer A. The solutions were vigorously vortexed before adding onto cells. NanoBRET was measured in a similar manner to BRET as described above, except the luminescence and resonance were measured at 447 nm and 610 nm respectively. The ratios and values were calculated as below

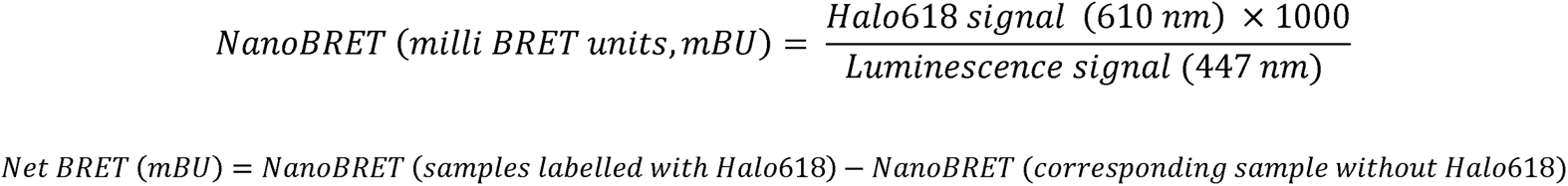

### Nanoluciferase complementation based SPASM assays

Cells expressing the T2R14 linked to different CFTR domains through SPASM or block linker were stimulated with DPH (final concentration 2 mM, resuspended in buffer A) or with buffer A containing DMSO (0.4 %). The plate was centrifuged at 100x*g* for 30 seconds, cells were incubated for 5 minutes, with orbital shaking in a SpectraMax iD5 plate reader equilibrated to 37°C. Subsequently 25 µL NanoGlo Live cell substrate (diluted 1:500 in Live cell buffer) was added to each well, the plate was centrifuged as before, and returned to the plate reader. Following 30 seconds shaking, the luminescence signal was measured for 5 minutes, at every 1 minute interval with 500 ms integration per well. Total luminescence counts for the 5 minute interval were calculated as area under the curve (AUC calculated as area of multiple trapeziums). The AUC values of buffer treated samples were subtracted from the values of DPH stimulated samples, and the difference was normalized to cell surface expression of the specific sensor (see immunofluoresce below). This final value is represented as the Δ luminescence.

### Immunofluorescence

HEK293T cells expressing SPASM or block domains were plated at 60000 cells per well in poly-L-lysine coated black plate with transparent bottom. Following adherence for 4 hr, the cells were washed with 1x PBS and fixed with 2.5% paraformaldehyde for 15 min. Fixed cells were not permeabilized but labeled with anti-HA primary antibody (1:100) in blocking solution (PBS containing 2 mg/mL BSA) for 30 minutes, washed thrice with 1x PBS, labeled with AlexaFluor488-tagged anti-mouse secondary antibody (1:500) for a further 30 minutes. Unbound secondary antibody was removed by washing with 1x PBS. In parallel, non-specific signal was determined from cells expressing the same SPASM sensor, but not treated with the primary antibody. Fluorescence (excitation 470 nm, emission 505 nm) was measured in an iD5 plate reader. Sensor expression was determined by subtracting the value of the ‘no primary control’ from the fluorescence of samples treated with the anti-HA primary antibody. This value was used to divide the subtracted AUC values to obtain the Δ luminescence values.

### Data analysis and statistics

Data acquired in the plate reader was analyzed using Microsoft Excel, and graphs and statistical analysis were performed using GraphPad Prism.

## RESULTS

### T2R14 and CFTR interact with each other

BRET has been used to investigate protein-protein interactions (PPIs) involving GPCRs in a cellular context [42]. We could detect BRET between the donor-acceptor pair formed by Nanoluciferase and EYFP, when the proteins were in proximity, but not when they were separated by a rigid ‘block’ domain derived from TRAF2, and recommended previously [40] (**Supplementary figure 1A**). Following subtraction of the Nanoluciferase signal, net BRET was found to decrease dramatically upon separation of EYFP and Nanoluciferase by the rigid ‘block’. (**Supplementary figure 1B**). Having validated the donor-acceptor pair, hardware and experimental parameters, we then used the BRET assay to query the direct PPIs between a known interacting pair of transmembrane proteins. T2R14 and β2-AR are known to interact with each other, with the latter serving as a chaperone for the former [43]. T2R14-Nanoluciferase and β2-AR-EYFP were co-transfected into cells at increasing ratio of acceptor plasmid DNA (β2-AR-EYFP) compared to the donor (T2R14-Nanoluciferase). BRET ratios and net BRET values increased with increasing acceptor and reached saturation (**Supplementary figure 1C**). Thus, the specific interaction between two transmembrane proteins, T2R14 and β2-AR, could be detected using BRET between Nanoluciferase and EYFP. We therefore used Nanoluciferase and EYFP as the BRET pair to investigate the possibility that T2R14 and CFTR interact with each other.

T2R14 tagged with a C-terminus Nanoluciferase (T2R14-NLuc) served as the donor (D) and CFTR with EYFP at the N-terminus (EYFP-CFTR) was used as the acceptor (A) (**Figure 1A**). HEK293T cells were transfected with either T2R14-NLuc + EYFP-CFTR or T2R14-NLuc alone. BRET ratio was measured from these cells following treatment with either the T2R14 agonist DPH (2 mM, corresponding to E_Max_) [44], or with 0.4% DMSO (**Figure 1B**). The net BRET was calculated in the DPH treated and buffer treated conditions, and found to be higher upon DPH stimulation (p<0.05). The increase in net BRET indicates that DPH-treatment causes or permits T2R14 and CFTR to interact with each other at close proximity. Further, net BRET in DPH-treated cells increases with increasing ratio of EYFP-CFTR (acceptor) to T2R14-NLuc (donor) (**Figure 1C**), indicating interactions between these proteins. Saturation of BRET ratio at higher acceptor:donor ratios is a characteristic of specific PPIs [45]. Our efforts to achieve acceptor:donor ratio greater than 4:1 were stymied by the toxicity of transient transfection at higher DNA amounts and the minimum amount of T2R14-NLuc plasmid required to observe Nanoluciferase activity. For comparison, we examined BRET between β2-AR-NLuc and EYFP-CFTR at increasing CFTR: β2-AR ratio. The net BRET ratio remained unchanged between β2-AR-NLuc (donor alone) and co-transfection of β2-AR-NLuc and EYFP-CFTR, even at increasing CFTR ratio, indicating that CFTR and β2-AR did not interact directly with each other, consistent with previous observations (**Supplementary figure 1D**) [46]. Thus, DPH-stimulated T2R14 and CFTR interact specifically and directly with each other.

**Figure 1:**
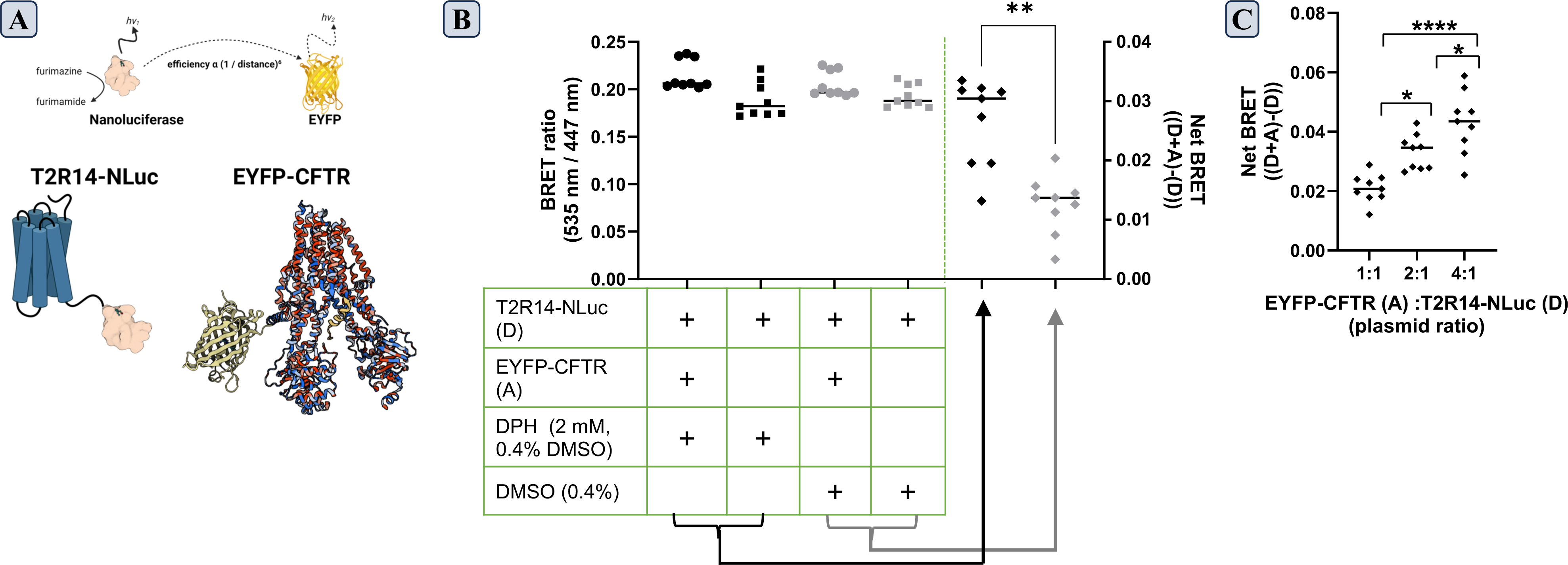
Agonist-dependent interaction between T2R14 and CFTR. **(A)** Schematic representation of Bioluminescence Resonance Energy Transfer (BRET), a non-radiative process such that BRET efficiency is inversely proportional to the sixth power of the distance between the donor (Nanoluciferase) and acceptor (EYFP). To test possible interaction, T2R14 was tagged with Nanoluciferase (T2R14-NLuc, donor (D)) and CFTR with EYFP (CFTR-EYFP, acceptor (A)), shown to scale. **(B)** Cells transfected with plasmids encoding either donor (D) or donor + acceptor (D+A) were treated with either DPH (2 mM) or DMSO (as control) for 5 minutes, and the luminescence (447 nm) and resonant (535 nm) signals were recorded and converted to BRET ratio (535 nm / 447 nm), represented on the left Y-axis. For each treatment (DPH / DMSO), the net BRET ratio was calculated as (BRET ratio (D+A) – BRET ratio (D)) and plotted on the right Y-axis. Net BRET ratio is different in DPH-treated cells compared to DMSO-treated controls (** indicates p <0.01, paired *t-*test, 9 observations from 3 biological replicates, median value indicated by horizontal bar). **(C)** Net BRET in DPH-treated cells co-transfected with the specified ratio of donor and acceptor plasmids. Net BRET increases with an increase in acceptor:donor plasmid. Graph displays data from 9 observations across 3 biological replicates, analyzed by one-way ANOVA and further tested by Tukey’s post-hoc test. ** indicates differences are significant at p<0.05, **** - differences are significant at p<0.0001. Median value indicated by horizontal bar.

### N- and C-terminus domains of CFTR interact with T2R14

We subsequently sought to dissect the molecular determinants of interaction between T2R14 and CFTR. For this purpose, a protein engineering-based approach, termed Systematic Protein Affinity Strength Modulation (SPASM) that has been used to investigate various protein-protein interactions [47] was employed. CFTR is a multi-domain protein with various cytoplasmic domains connected through twelve transmembrane domains [48] **(Figure 2A)**. We reasoned that one of the cytoplasmic domains of CFTR is likely to interact with the cytosolic interface of T2R14. Hence, SPASM sensors were constructed linking T2R14 to individual cytoplasmic domains of CFTR through a stable single alpha helix (10 nm ER/K linker) with split Nanoluciferase domains - LgBiT and SmBiT [49] - at opposite ends (**Figure 2A**). It was hypothesized that upon interaction between T2R14 with the appropriate CFTR domain, LgBiT and SmBiT will come in proximity, allowing for Nanoluciferase complementation. Therefore, an increase in luminescence from cells expressing a particular SPASM sensor would indicate the interaction of that specific CFTR domain with T2R14 **(Figure 2B)**. This split Nanoluciferase design of the SPASM sensor was tested by replacing the fluorophores in the well-characterized SPASM sensor β2-AR-Spep, with LgBiT and SmBit domains of Nanoluciferase (**Supplementary figure 2A**). Luminescence was measured from cells transfected with the modified β2-AR-Spep and treated with increasing concentrations of isoproterenol (10 pM to 1 mM). Increasing the isoproterenol concentration led to an increase in luminescence, which saturated at higher concentrations of isoproterenol, and exhibited EC_50_ ∼100 nM (**Supplementary figure 2B**). Thus, the split Nanoluciferase complementation is an appropriate system to report PPIs through SPASM sensors.

**Figure 2:**
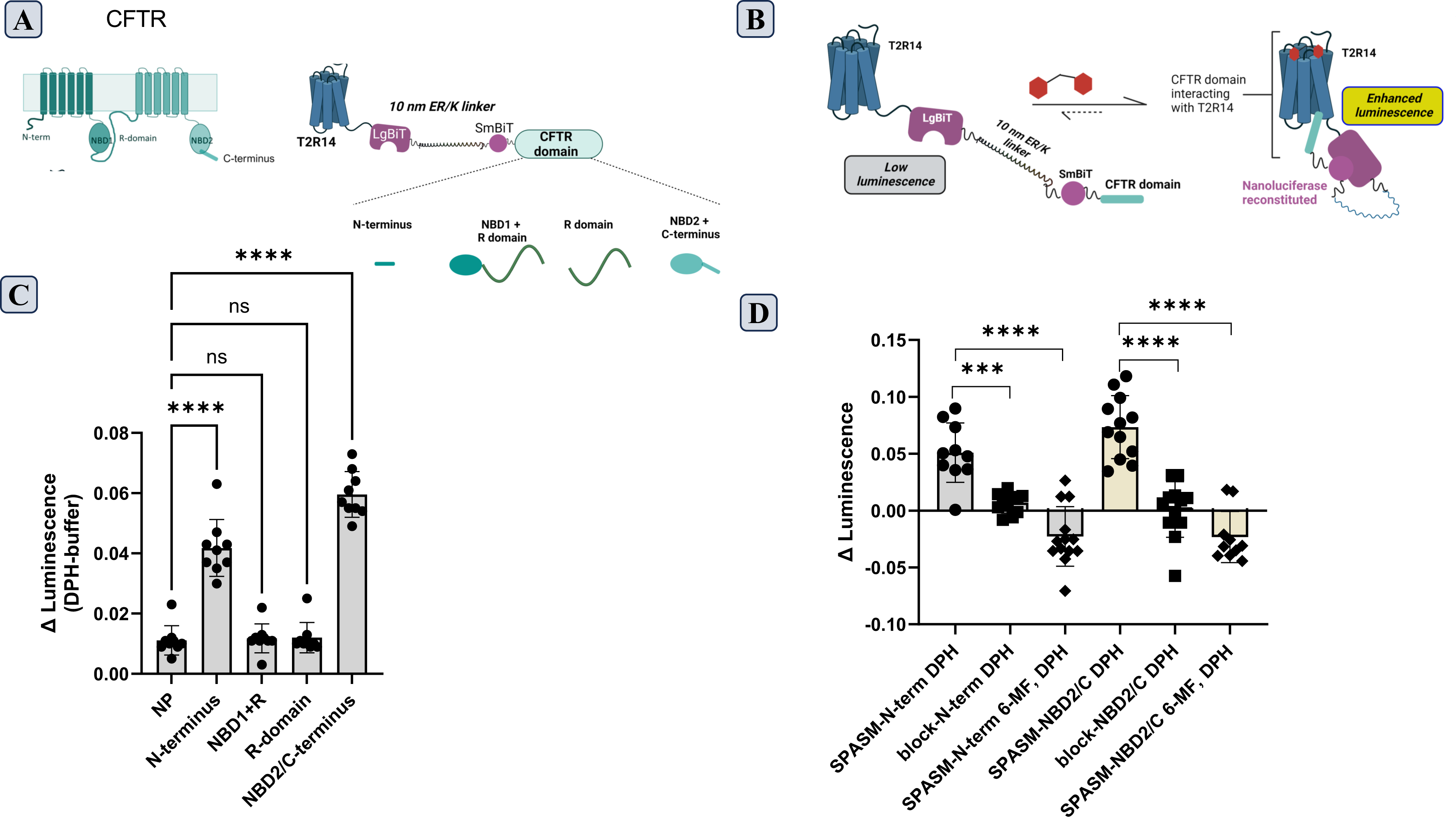
Interaction between intracellular domains of CFTR and T2R14. **(A)** Schematic representation of CFTR highlighting the intracellular domains (left) and SPASM sensors linking T2R14 to each of the specified CFTR domains through a core sensor module consisting of LgBiT, 10 nm ER/K linker, SmBiT made contiguous by the insertion of (GSS)_4_ flexible linker (right). **(B)** Principle of SPASM sensor, wherein interaction between CFTR domain and T2R14 results in Nanoluciferase complementation detected by an increase in luminescence. **(C)** HEK293T cells transfected with the indicated SPASM sensors linking T2R14 to a specified domain of CFTR were stimulated with 2 mM DPH or treated with buffer. Luminescence was measured for 5 minutes following the addition of NanoGlo Live Cell substrate®, and area under the curve (luminescence vs time) was calculated for each sensor. Difference between the luminescence from DPH stimulated vs buffer treated conditions, normalized to sensor expression, is represented as Δ luminescence. Data from 9 observations across three biological replicates analyzed by one-way ANOVA and further tested by Tukey’s post-hoc test. **** - indicates differences are significant at p<0.0001. **(D)** HEK293T cells transfected with either SPASM or ‘block’ sensors linked to either CFTR N-terminus or CFTR NBD2/C-terminus, were pre-treated with 6-methoxyflavonone (6-MF, 500 µM) or buffer; followed by stimulation with 2 mM DPH or buffer. Δ luminescence value is calculated as described in (c) above. Data from at least 10 observations across four biological replicates analyzed by one-way ANOVA and further tested by Tukey’s post-hoc test. *** indicates differences are significant at p<0.002, **** - differences are significant at p<0.0001. Mean value indicated by height of the bar.

Having validated the split Nanoluciferase complementation module in the context of SPASM sensors, we investigated the possibility of interaction between T2R14 and different domains of CFTR, using the corresponding sensors. PPI between T2R14 and CFTR domains were evaluated in the absence and presence of the bitter agonist DPH. Two sensors, T2R14-N-terminus and T2R14-NBD2/C-terminus, that linked T2R14 to either CFTR N-terminus lasso motif (aa 1 - 80) [50] or to the CFTR NBD2-C terminus (1151-1480) respectively, exhibited a DPH-dependent increase in luminescence, which was significantly greater than the change in luminescence observed with the control sensor (NP) harboring no CFTR domain (**Figure 2C**). The other sensors linking T2R14 to either the NBD1 and R domain or to the R domain alone did not exhibit any change in luminescence relative to the control sensor. The increase in luminescence observed with the T2R14-SPASM-N-terminus and T2R14-SPASM-NBD2/C-terminus sensors suggests that DPH-stimulated T2R14 is capable of interacting with either the CFTR N-terminus lasso motif, or with the CFTR NBD2/C-terminus, independently. Further, we replaced the ER/K linker domain in T2R14-N-terminus and T2R14-NBD2/C-terminus SPASM sensors with the rigid ‘block’ domain used previously. Introduction of the ‘block’ domain prevented the DPH-stimulated increase in luminescence (**Figure 2D**). Interaction between T2R14 and the identified domains of CFTR is contingent on the presence of a permissive linker. In the T2R14-N-terminus and T2R14-NBD2/C-terminus sensors, pre-treatment of cells with 6-methoxy flavanone (6-MF), a known antagonist of T2R14, prevented the subsequent DPH-dependent increase in luminescence. Thus, T2R14 – CFTR interactions are a specific property of the N and C-terminus cytoplasmic domains of CFTR, and are limited to the DPH-stimulated state of T2R14.

### T2R14-G protein interactions in response to microbial signals

Similar to DPH, bacterial compounds stimulate T2R14, which in turn leads to antimicrobial responses [10–12]. Therefore, we asked if QSMs from CF-related microbes could affect T2R14-CFTR interactions. Along with AHL-C12 which is a QSM from *P. aeruginosa*, and has been well-characterized as a T2R14 agonist [10, 11], we also tested the fungal quorum sensing compounds – farnesol [51] and tyrosol [52], which have been found to stimulate innate immune responses downstream of T2R14 (Singh et al., manuscript under revision). We employed NanoBRET (**Figure 3A**) [53], to determine the effect of these selected QSMs on T2R14-G protein interaction, with T2R14-Nanoluciferase as the donor and Gα subunit-HaloTag as the acceptor. The appropriate acceptor orientation for each G protein isoform was chosen based on DPH stimulated changes in NanoBRET signal (**Supplementary figure 3,** and **Singh et al.,** manuscript under revision). DPH-stimulation of cells expressing T2R14-NLuc and Gαi-HaloTag led to increased NanoBRET signal, consistent with the known signaling pathway downstream of agonist-stimulated T2R14 [14, 15, 54]. Interestingly, DPH-stimulated T2R14-NLuc also exhibited an increase in NanoBRET with Gαq-HaloTag signifying interaction between T2R14 and Gαq (**Figure 3B**). Therefore, the ability of microbial compounds to mediate coupling with Gαi and Gαq was tested. Similar to DPH, treatment of cells with either AHL-C12 or farnesol or tyrosol resulted in an increase in net NanoBRET values (**Figure 3C, D**). Thus, all three QSMs from CF-relevant microbes stimulated coupling of T2R14 to both Gαi and Gαq. This suggests that microbial quorum sensing molecules may signal through multiple G protein pathways downstream of T2R14.

**Figure 3:**
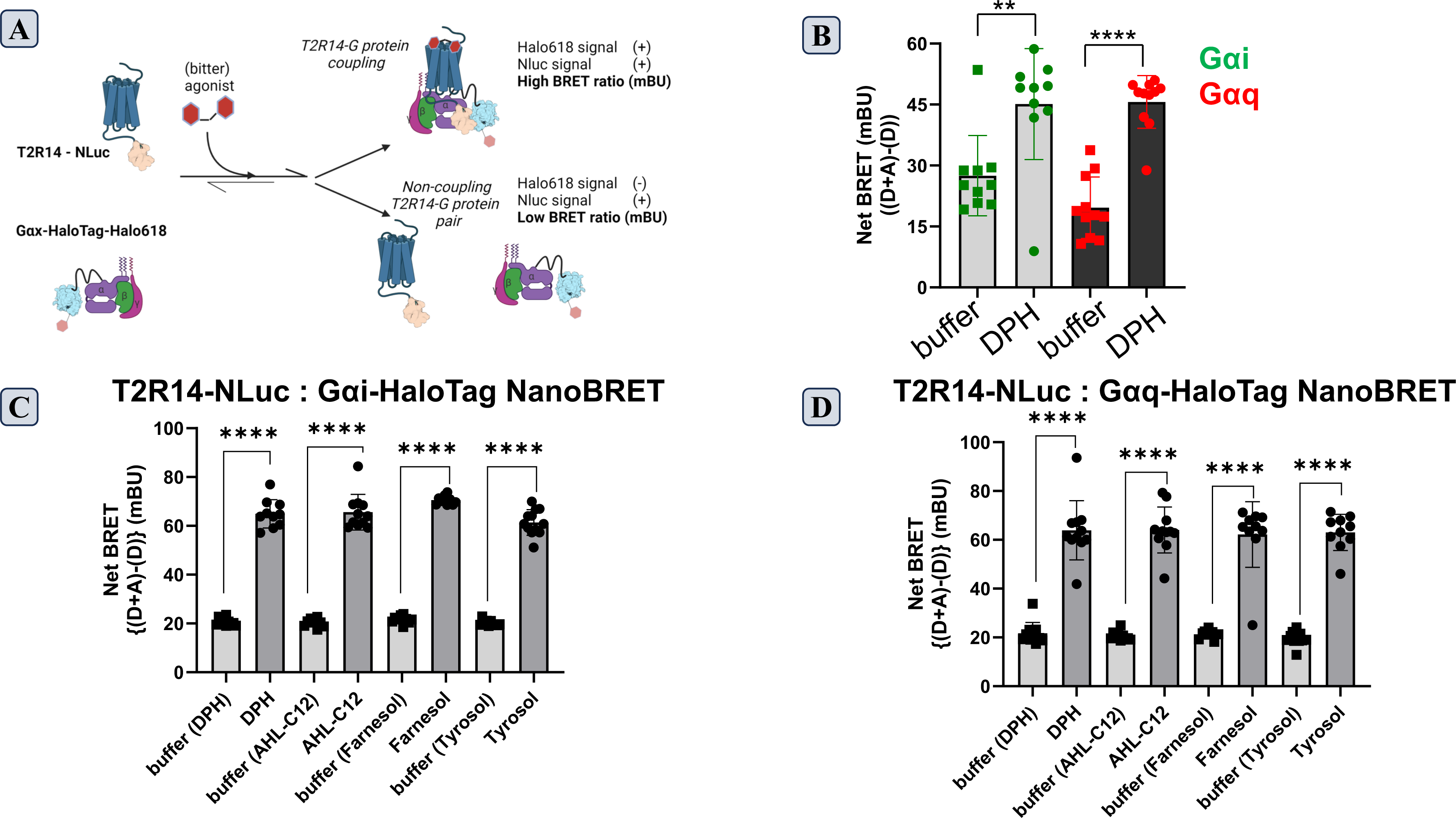
Multiplicity of G protein signaling downstream of T2R14. **(A)** Schematic of the NanoBRET assay to detect T2R14 – G protein interactions. The optimized orientations involve T2R14-Nanoluciferase as the donor (D) and Gα subunits with HaloTag labelled with Halo618, as the acceptor (A). T2R14-G protein coupling in response to ligand-stimulation results in an increase in BRET ratio between Acceptor (610 nm) and Donor (447 nm) ratio. **(B)** NanoBRET was measured (610nm / 447 nm) in response to DPH stimulation of HEK293T cells co-transfected with T2R14-NLuc and Gαi-HaloTag or T2R14-NLuc and Gαq-HaloTag. The difference between BRET ratio in the presence or absence of Halo618 is Net BRET. Net BRET increases in response to DPH (2 mM), for both pairs investigated. **(C, D)** Net BRET measured in HEK293T cells transfected with T2R14-NLuc and Gαi-HaloTag (C) or T2R14-NLuc and Gαq-HaloTag (D), and stimulated with microbial products - AHL-C12, farnesol or tyrosol (200 µM each); or corresponding buffers/solvent control. Net BRET increases in response to all three ligands, in T2R14-Gi as well as T2R14-Gq pairs. Graphs contain data from at least 10 observations across 4 biological replicates, analyzed by unpaired t-test, ** indicates differences are significant at p<0.05, **** - differences are significant at p<0.0001. Bar height represents the mean value.

## DISCUSSION

The human genome encodes at least 25 bitter taste receptor genes, that are expressed not just in gustatory context but also in extra-oral tissues [5, 37, 55]. T2R14, which is expressed in bronchial epithelia, senses microbial QSMs and responds by eliciting components of the innate immune response [10, 11]. Considering the tissue context (bronchial epithelium) of cystic fibrosis, the occurrence of opportunistic respiratory infections in pwCF, the high expression levels of T2R14 in this tissue and its role in antimicrobial defense, we considered whether T2R14 could contribute to the removal of microbes and affect outcomes in pwCF. Cystic fibrosis is caused by mutations in the *CFTR* gene encoding the cystic fibrosis transmembrane conductance regulator [22, 23], an anion channel that helps in the secretion of mucus, which traps bacteria and expels them from the respiratory tract. There is accumulating evidence that activities of bitter taste receptors and of ion channels and transporters are mutually affected [34–36]. In the context of the respiratory epithelium, CFTR and T2Rs are reported to affect each other but any direct protein-protein interactions have not been reported. Further, CFTR is reported to interact with different GPCRs, either directly or indirectly. Hence, we investigated the possibility that CFTR and bitter taste receptors, specifically T2R14, interact with each other.

BRET is a biophysical technique that reports protein-protein interactions with close proximity and a resolution of approximately 10 nm. We found that BRET between T2R14 and CFTR increased specifically upon application of a T2R14-agonist (**Figure 1**), indicating that under a bitter tastant stimulus to T2R14, CFTR and T2R14 interact with each other. A similar approach, FRET (Förster Resonance Energy Transfer) has been previously applied to detect CFTR interaction with Syntaxin-17 [38]. Relocalization of proteins in response to a stimulus facilitates interactions between partner proteins in a complex [56]. However, HEK293T cells used here are not grown in conditions that would allow for the formation of motile cilia or apico-basal polarity, suggesting that CFTR and T2R14 are not spatially segregated. Hence agonist-stimulated relocalization is unlikely to drive CFTR-T2R14 interaction. However, conformational selection may govern the CFTR-T2R14 interaction, such that DPH-bound T2R14 adopts a conformation that can interact with CFTR.

To further dissect T2R14 – CFTR interaction, we used SPASM, a protein engineering approach that tethers two candidate protein entities through a stable single alpha-helix (SAH) of known length (10 nm) composed of repeating acidic and basic residues (E_4_(R/K)_4_) [39], and a bi-partite reporter system -composed in this case of the large and small parts of Nanoluciferase, LgBiT and SmBiT respectively. The SAH domain ensures stoichiometric expression of the proteins, and increases the effective concentration of PPI, without enforcing the interaction by forced proximity. SPASM sensors have been previously used to investigate the details of inter- and intra- molecular PPIs through FRET [47]. Here, we have replaced the FRET reporter module with a Nanoluciferase complementation system such that upon interaction between the candidate proteins, LgBiT and SmBit are brought in proximity and Nanoluciferase activity is reconstituted. From the CFTR side, the N-terminus lasso motif and the NBD2/C-terminus domains were detected to interact with T2R14 in response to DPH, using the luminescence-based SPASM assay (**Figure 2**). The NBD1 and Regulatory (R) domain-containing SPASM sensors did not exhibit T2R14 interactions. CFTR has previously been reported to interact with other transmembrane proteins including SNARES, Syntaxin1A [57] and Syntaxin17 [38]. The Syntaxin1A interaction has been localized to the N-terminus of CFTR, while Syntaxin17 predominantly interacts with the N-terminus, but also exhibits interactions with the NBD1 and NBD2/C-terminus. The observations from SPASM approach employed in this work, which reveal that T2R14 is capable of interacting with either the N-terminus lasso motif or the NBD2/C-terminus of CFTR, are supported by these precedents.

The agonist-dependent interaction between T2R14 and CFTR is also supported by the evidence that pretreatment with 6-methoxy flavonone, the T2R14 antagonist [58], prevents the interaction with CFTR domains. The CFTR interactome has been profiled under various conditions, including mutations in CFTR, drug-treated conditions; and using multifarious technical approaches. The following GPCRs have been identified as direct CFTR interactors - GPRC5A, M4R, 5HT5AR, MC1R, MC3R, P2RY6, VIPR-1 and a few odorant receptors [59–61]. CFTR is also reported to interact via the PDZ-domain and linker proteins with β2-AR [46] and A2A-R [62]. These -omics or high throughput approaches were designed from a CFTR perspective, rather than a GPCR perspective. Therefore GPCR-agonist dependent interactions with CFTR were possibly not selected for. The results from this work add T2R14 as another interactor of CFTR.

The N-terminus lasso motif of CFTR is analogous to the lasso loop in other ABC transporters such as Ycf1 [63] and MRP1 [64]. In these proteins, the lasso loop interacts with the NBD1 and R-domain, particularly when NBD1 is phosphorylated. Such structural stabilization in context of CFTR could reduce the activity. The agonist-dependent interaction between T2R14 and CFTR therefore leads to a possible mechanism whereby T2R14 interacts with the lasso motif and sequesters it away from CFTR, enhancing CFTR activity. The ability of microbial QSMs to selectively stimulate the T2R14-CFTR interaction suggests that CFTR activity could be enhanced in response to a physiologically relevant stimulus, through the agency of T2R14. Such an increase in CFTR activity could contribute to mucociliary clearance and improve expulsion of microbes from the respiratory tract. The significance of NBD2 interaction with T2R14 is not obvious to us at this time.

A key insight from this work is the multiplicity of G protein signaling downstream of agonist-stimulated T2R14, beyond the canonical Gi-pathway characterized in extraoral contexts (**Figure 3**). Early efforts to understand bitter taste receptor signaling used the promiscuous subunit Gα16 fused to differing lengths from the C-terminus of gustducin and also identified Gαi2 as the other cognate G protein [14]. A reconstitution approach to investigate T2R – G protein coupling did not detect interaction between membranes containing mT2R5 and Gq or Gs [15]. The NanoBRET approach employed here utilizes full-length non-chimeric G proteins and is executed in live cells to reveal that agonist-stimulated T2R14 interacts with Gαq, in addition to the expected Gαi interaction. Previous work has found Ca^2+^ signaling downstream of T2R14, and attributed it to the βγ arm of Gi signaling. However, recent reports suggest that GPCR-mediated Ca^2+^ signaling, previously attributed to the βγ arm, requires Gαq activation [21]. The observation here that T2R14 stimulation leads to engagement of Gαq (in addition to Gαi), therefore provides a comprehensive mechanism that explains with multiple previous observations that T2Rs stimulation results in a Ca^2+^ spike, and is consistent with more recent reports. The NanoBRET assays were not designed to detect basal activities or agonist-free T2R14-G protein association or precoupling, such as was reported in context of the recent cryoEM structure of T2R14 [20].

The impact of CFTR domains on T2R14 coupling to Gi versus Gq, and the effect of CFTR domain-T2R14 interactions on downstream signaling pathways are being studied. These efforts will examine the biased signaling downstream of T2R14 in response to the selected QSMs, and its modulation by the CFTR domains. The work here identifies the N- and C-terminus domains of CFTR interact with T2R14. Future efforts could delve into fine mapping this interaction at the CFTR interface(s), and identifying and mapping the interface from T2R14. The effect of T2R14 on CFTR activity warrants separate investigation, particularly in the context of various CF-causing mutants. There may be translational value in examining the interplay between T2R14 signaling, CF-causing mutants and highly effective modulator therapy (HEMT).

## ACKNOWLEDGEMENTS

Schematics and illustrations were created with BioRender.com.

This work was supported by a research grant from the Cystic Fibrosis Foundation (003100G221-Chelikani).

---

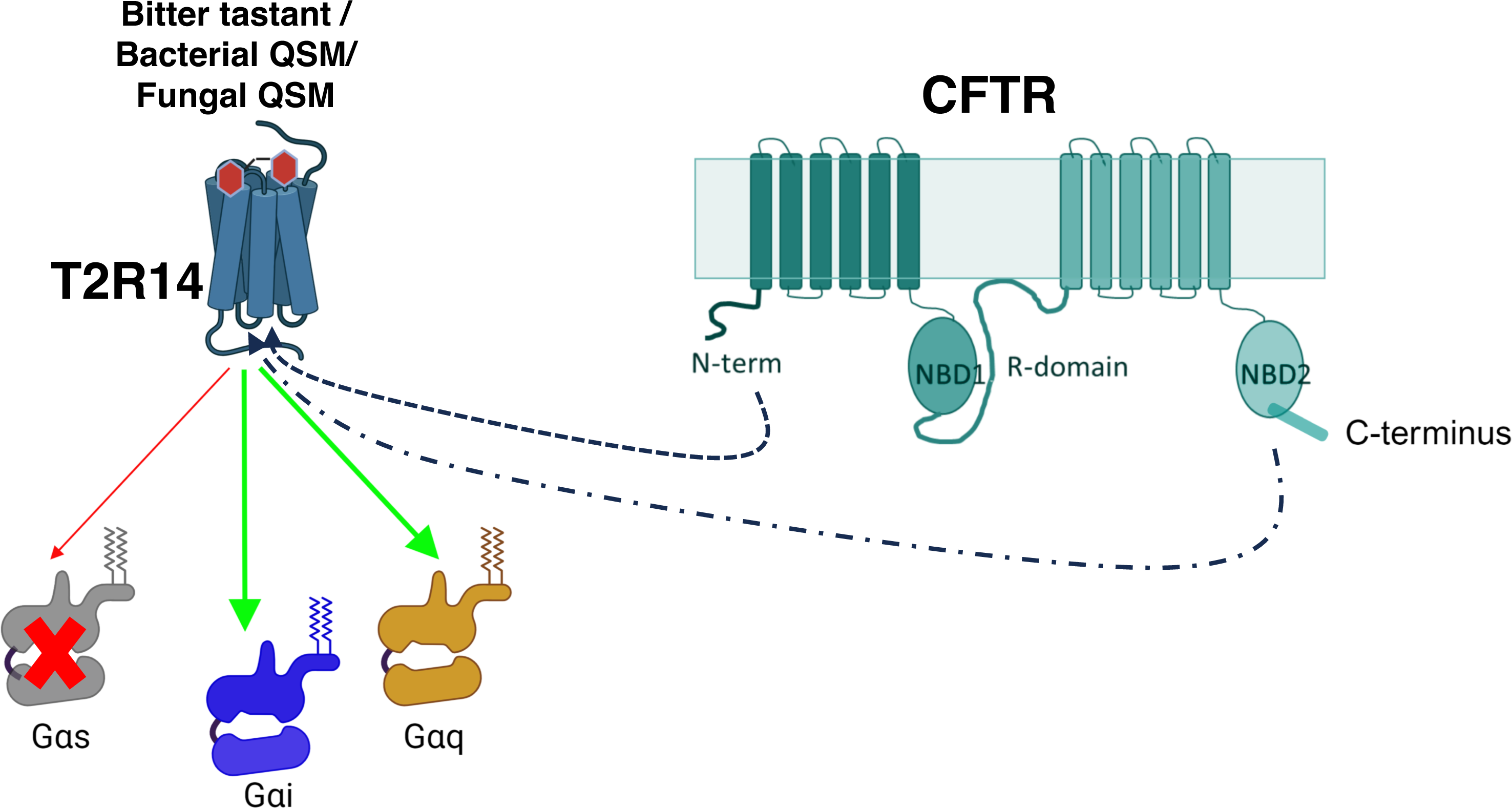
Graphical Abstract.

