## Supplementary Figures 1-3 for "T2R14 mediated antimicrobial responses through interactions with CFTR"

**Supplementary Figure 1:** Validation of BRET measurements

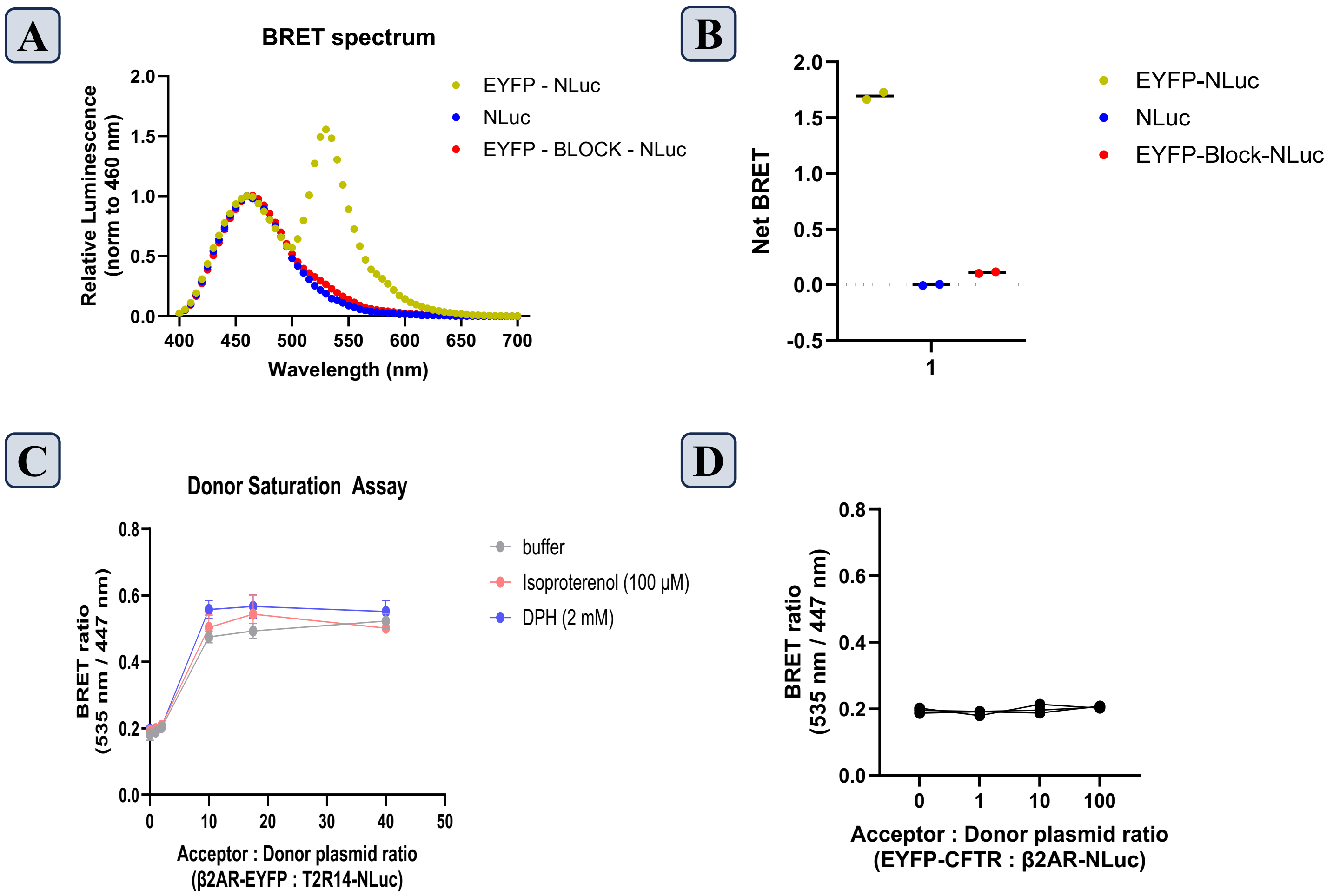

**Supplementary Figure 2:** Validation of SPASM sensor based on split Nanoluciferase complementation

A

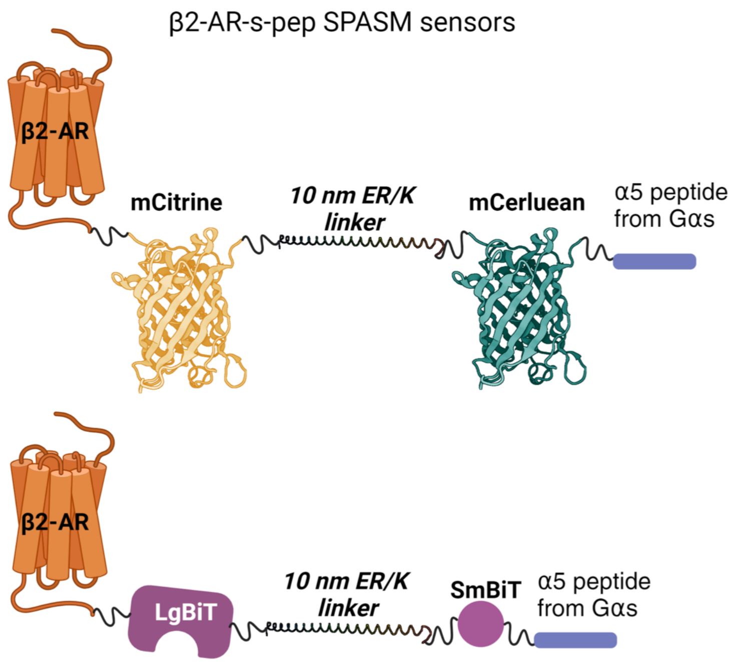


B

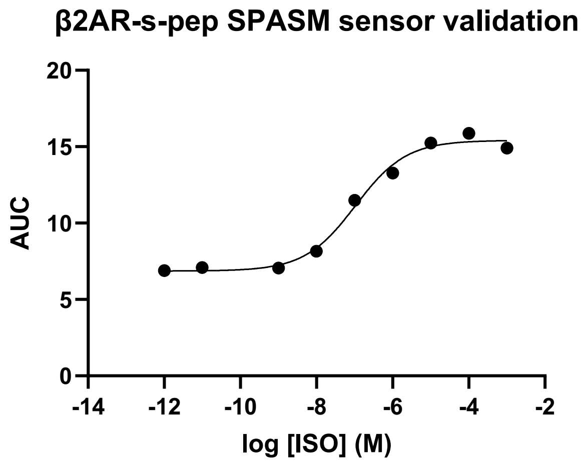

**Supplementary Figure 3:** Determination of probe orientations for NanoBRET

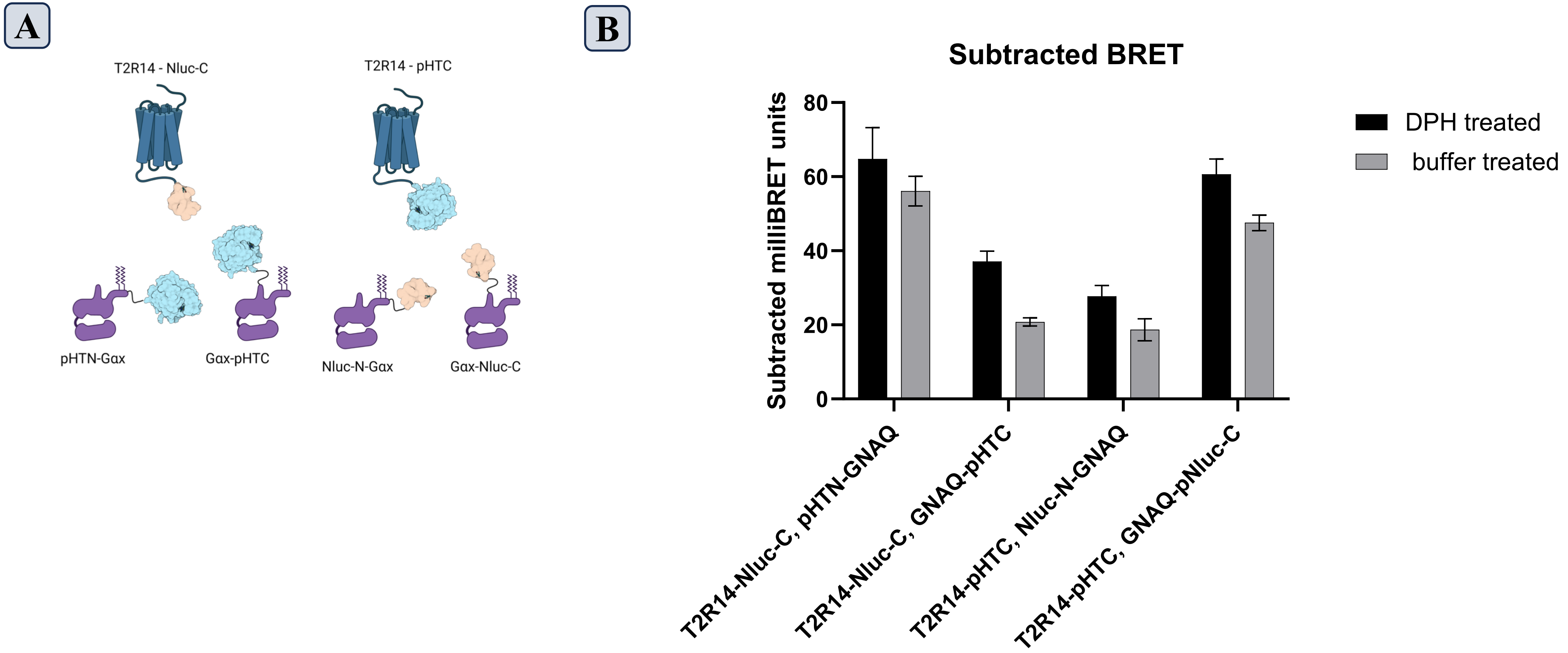
